# Figuring things out: Innovation arena in wolves and dogs

**DOI:** 10.64898/2026.10.26.761969

**Authors:** Dániel Rivas-Blanco, Femke Van Steen, Sabine Tebbich, Sarah Marshall-Pescini, Friederike Range

## Abstract

Dogs and wolves present an interesting lens through which to examine innovative behaviors, as both of their environments promote the development of innovations for different reasons. Dogs’ natural environment (as free-ranging animals) is characterized by high diversity and abundance in novel feeding opportunities. Conversely, wolves live in an unpredictable and harsh environment in which adapting to sudden scarcity is vital for survival. Thus, to analyze the role of the environment in innovative behavior (both evolutionarily and ontogenetically) we studied problem-solving skills of three populations: captive wolves, “pack dogs” (raised similarly to the wolves), and pet dogs. For this, we used an “innovation arena” (after Rössler & Mioduszewska et al., 2020) consisting of 9 novel puzzle boxes presented simultaneously. Pet dogs were better at solving the boxes than their pack-living counterparts, which could point towards prior positive experiences with similar opportunities (here, human-created constructs) providing motivation to interact with novel features of the environment, which would eventually lead to innovation. We found little effect of evolution: wolves needed more time to approach the boxes (potentially due to neophobia), but both wolves and pack dogs interacted with and solved the boxes to a similar extent.

## Introduction

In an ever-changing world, animals are constantly presented with both novel challenges and novel opportunities. Innovation, defined as the discovery of new behavioral interactions with either the physical or social environment (Tebbich et al., 2016), constitutes a valuable tool for animals to adapt to their environment to deal with these novel features.

The innovation process is complex, with many traits influencing the emergence of novel behaviors and the likelihood of being adopted into an animal’s regular behavioral repertoire (Chow et al., 2016, Griffin & Guez, 2014). For instance, innovation is inhibited by neophobia, defined as an animal’s fear towards novel features of the environment. Conversely, an animal’s drive to approach novel objects and engage in novel situations (neophilia), and their willingness to interact with these novel opportunities for extended periods of time (persistence of exploration) are traits that promote innovation (Chow et al., 2016, Griffin & Guez, 2014).

An animal’s ecology also has a great impact on their likelihood to engage in innovative behavior. Generalist species, for example, are more likely to develop innovations than specialist species, both because they are more likely to encounter novel features of their environment they can exploit and because they would profit the most out of novel feeding opportunities (Ducatez et al., 2015; Tebbich et al., 2016).

However, regardless of an animal’s traits, innovation is largely dependent on chance, as the individual needs to come across a novel opportunity to begin with. It would follow then that particularly complex and heterogeneous environments, rich in resources that animal could exploit would select for traits that promote innovation (Tebbich et al., 2016, Greenberg & Mettke-Hoffman, 2001). This would also be the case for unpredictable and harsh environments, in which resources may be scarce temporarily or permanently (Tebbich et al., 2016, Greenberg & Mettke-Hoffman, 2001).

Interestingly, wolves and dogs’ evolutionary history provides a peculiar lens through which to explore the selective pressures underlying innovation. Both species are generalists and therefore could greatly benefit from exploiting novel feeding opportunities. However, dogs’ environment is far richer in such opportunities, as they inhabit human-shaped landscapes, which are characterized by a diverse environment, abundant in novel opportunities (Griffin et al., 2017). This might also have been relevant during dogs’ domestication process, as “early” dogs needed to adapt to the human social and ecological niche (Coppinger & Coppinger, 2002); forming bonds with humans and living as scavengers relying on anthropogenic food sources (Butler & Du Toit, 2002; Newsome at al., 2014). Alternatively, however, resources such as human feces, which remains an important source of nutrients for modern-day free-ranging dogs (Butler et al., 2018; Butler & Du Toit, 2002), were usually abundant and easy to access in the last 10 000 years of the domestication process. In such a rich environment, selective pressures towards innovative problem solving may have been relaxed in dogs (similar to what was discussed by Frank, 1980). This is contrasted by wolves, who usually inhabit environments less prone to rapid change, but rely on food sources that are unpredictable and infrequent (Mech et al., 2015). Therefore, although dogs live in an environment which offers many opportunities to access new resources, they may not necessarily have the need to do so. Conversely, wolves would have access to far fewer such opportunities, but as opposed to dogs, learning to exploit them could be more important for survival.

Additionally, dogs offer the opportunity of examining the role of experience since, depending on the population, they differ in the extent of contact with human-shaped environments. For example, pet dogs usually have more experience with novel opportunities arising from human-related activities than dogs living in kennels. As discussed by Tebbich et al. (2016), prior experience with features of the environment plays a crucial role in innovative behavior, as it equips animals with the ability to exploit “adjacent opportunities”; novel features of the environment similar to others that the animal has had a priori history of reinforcement with, and that can be exploited through generalization.

Thus, to test the innovation abilities of two populations of dogs (one comprised by pets, and the other of pack-living animals) and one population of wolves (raised in a similar manner as the pack-living dogs), we used an adapted version of Rössler & Mioduszewska et al. (2020)’s innovation arena. They developed this paradigm to test the innovative problem-solving skills of wild and captive Goffin cockatoos. The main feature of this setup is that it is comprised of several puzzle boxes with different opening mechanisms that are presented simultaneously to the animals. Through this design, animals are provided with a multitude of opportunities to engage in problem solving, as the opening mechanisms of each of the boxes constitutes a unique problem that needs to be solved independently, thus increasing the chances for innovative problem solving, even after subsequent presentations of the same apparatuses. Rössler & Mioduszewska found that factors related to general task engagement (e.g., their interaction with each one of the apparatuses, and the time spent in their vicinity) and the number of apparatuses solved relative to the ones they interacted with were the main predictors of problem-solving success.

The purpose of the study was to study differences in innovative problem-solving between wolves and dogs through the framework of the innovation arena. More specifically, the aims of this study were to 1) map the different ways the animals interact with the novel puzzle boxes, 2) determine whether these different patterns of interaction may come as a result of evolution (wolves vs. pack dogs) or ontogeny (pack dogs vs. pet dogs), and 3) analyze how these patterns of interaction influence an animal’s likelihood to solve the puzzle boxes.

Marshall-Pescini et al. (2017a)’s social ecology hypothesis postulates that the differences in behavior between wolves and dogs are a result of adaptations to their environment, both in what pertains to their respective habitats and social structures. As such, we expected pack dogs to show a higher level of engagement with the apparatuses than wolves due to reduced neophobia and increased selection for their mainly scavenging lifestyle. However this would not necessarily translate into a higher rate of success, as they may lack the persistence of exploration required to solve the more complex apparatuses.. This would mirror their feeding patterns as scavengers in abundant and stable environments, where the optimal strategy is often to seek an alternative food source rather than trying persistently to reach for one that may be inaccessible (Brubaker et al., 2017; Rao & Bernasconi et al., 2018a). This would also match the performance of another canid species in a similar innovative problem-solving task, as foxes that lived closer to human-shaped environments did approach the apparatuses more frequently than their conspecifics from more rural areas, but did not have a greater rate of success (Morton et al., 2023).

We also expected pet dogs to be more explorative and potentially more successful than the pack-living ones. Pet dogs have more experience with human-made constructs in their daily lives, which should reduce their levels of neophobia, and could have also provided them with more knowledge on how to interact with some of the affordances present in our apparatuses through generalization. On top of that, since they have more positive experiences with human-created constructs (e.g., in the form of dog toys), their engagement with them should also be higher in general due to past reinforcement of explorative behavior by their owners (as discussed by Lazzaroni et al., 2019).

Wolves, however, have been shown to be more neophobic than dogs, potentially due to their evolutionary history of being hunted by humans (Range & Marshall-Pescini, 2022). However, while being more neophobic than dogs, wolves have also been shown to have more interest in novel objects, interacting with them for longer amounts of time than dogs (Moretti et al., 2015, Marshall-Pescini et al., 2017b, but see Rivas-Blanco & Gonnet-dit-Revel et al., 2025). This may be linked as well with their ecology, as they need to manipulate carcasses of their prey before eating them, a type of pre-feeding processing that has been linked with higher levels persistence of exploration (Mech et al., 2015; Mettke-Hoffman et al., 2002). Thus, we expected wolves to show a lower initial level of engagement and a higher latency to reach the apparatuses, but the engagement should increase and the latency decrease in later sessions. Additionally, previous studies (e.g., Rivas-Blanco & Gonnet-dit-Revel et al., 2025) have shown that some wolves are too neophobic to interact with novel objects at all. Accordingly, we predicted that the individuals that fail to interact with the apparatuses or do so in a very limited capacity should be mostly wolves.

## Methods

### Subjects

In this study, we tested pack-living wolves (*N*_wolf_ = 12; 8 males; mean age = 106.4 ± 9.9 months) and dogs (hereafter “pack dogs”; *N*_pack dog_ = 13; 5 males, mean age = 50.5 ± 12.6 months) at the Core Facility Wolf Science Center (WSC) in Ernstbrunn, Austria. Both the wolves and the pack dogs were hand-raised in similar conditions, with permanent contact to humans until the age of 5 months before being introduced into packs. They also lived under similar conditions after that point, in outdoor enclosures measuring 2000 to 10 000 m^2^.

In addition to these two populations, we tested pet dogs (*N*_pet dog_ = 9; 6 males, mean age = 102.0 ± 10.2 months). These pet dogs belonged to WSC trainers and lived in their houses as pets. We decided to recruit these dogs because they are genetically similar to the pack dog population (as several of the individuals were related to the pack dogs, see **Supplementary Table S1** for more details) and because they were already familiar with the testing environment that was used to test all three populations (something that has been shown to have an effect on dogs’ performance; Lazzaroni et al., 2020; Lazzaroni & Schär et al., 2023).

More information on the subjects that participated in this study can be found in **Table 1**. A more detailed account can be found in **Supplementary Table S1**.

**Table 1.** Subjects that participated in this experiment. The subject in gray (*Maikan*) was excluded from the experiment after showing an intense fear response after being exposed to all 9 apparatuses simultaneously in the first test session.

| Subject | Species | Population | Age at the start of the experiment (months) | Sex |
| --- | --- | --- | --- | --- |
| Amarok |  | Wolf | 112 | M |
| Chitto |  | Wolf | 111 | M |
| Etu |  | Wolf | 64 | M |
| Geronimo |  | Wolf | 146 | M |
| Kenai |  | Wolf | 136 | M |
| Maikan |  | Wolf | 64 | M |
| Nanuk |  | Wolf | 147 | M |
| Taima |  | Wolf | 64 | F |
| Tala |  | Wolf | 111 | F |
| Tekoa |  | Wolf | 64 | M |
| Una |  | Wolf | 112 | F |
| Yukon |  | Wolf | 146 | F |
| Enzi* | Dog | Pack | 84 | M |
| Hiari | Dog | Pack | 86 | M |
| Imara | Dog | Pack | 86 | F |
| Inaya | Dog | Pack | 11 | F |
| Kiano | Dog | Pack | 11 | M |
| Kovu | Dog | Pack | 14 | M |
| Layla | Dog | Pack | 118 | F |
| Moya | Dog | Pack | 9 | F |
| Taio** | Dog | Pack | 11 | M |
| Sarabi | Dog | Pack | 9 | F |
| Panya | Dog | Pack | 86 | F |
| Youma | Dog | Pack | 14 | F |
| Zuri | Dog | Pack | 118 | F |
| Asali | Dog | Pet | 127 | M |
| Bora | Dog | Pet | 118 | F |
| Freya | Dog | Pet | 72 | F |
| Hakima | Dog | Pet | 127 | M |
| Kilio | Dog | Pet | 136 | M |
| Koda*** | Dog | Pet | 121 | M |
| Pepeo | Dog | Pet | 85 | M |
| Zazu | Dog | Pet | 84 | M |
| Zhenga | Dog | Pet | 48 | F |

\*: Did not enter in the enclosure for session 4. He was given an additional session to compensate for that.
\*\* : Halfway through what would have been his 7th session, heavy rain started. Due to this, the session was cancelled and excluded from the analyses, and an additional session was conducted.
\*\*\*: For unknown reasons, the subject started showing an intense aversion towards the experiment after session 4, stopping interacting with the apparatuses and being hesitant about going inside the test enclosure. Due to this, all attempts to test him after the 4th session were excluded from data analyses.

### Apparatuses

The array of apparatuses was comprised of two sets of 9 puzzle boxes, one for the wolves (measuring 50×50×50 cm), and another for the dogs (measuring 20×20×20 cm). For each species, the 9 boxes were presented simultaneously in an arc to facilitate the movement of the subject from one to the next. Each apparatus was fixed to a squared table (50×50cm, 10 cm high for the dogs; 100×100cm, 20 cm high for the wolves) (see Figure 1). Each of the boxes had a different method to access the food reward, in the form of an “opening mechanism”. Most of these mechanisms were inspired by the study done by Rössler & Mioduszewska and colleagues (2020), albeit their number was reduced due to our species’ limited manipulation capabilities when compared with Goffin cockatoos. The position of the boxes was pseudo-randomized so that every box occupied a different position from the previous session for every subject.

**Figure 1.**
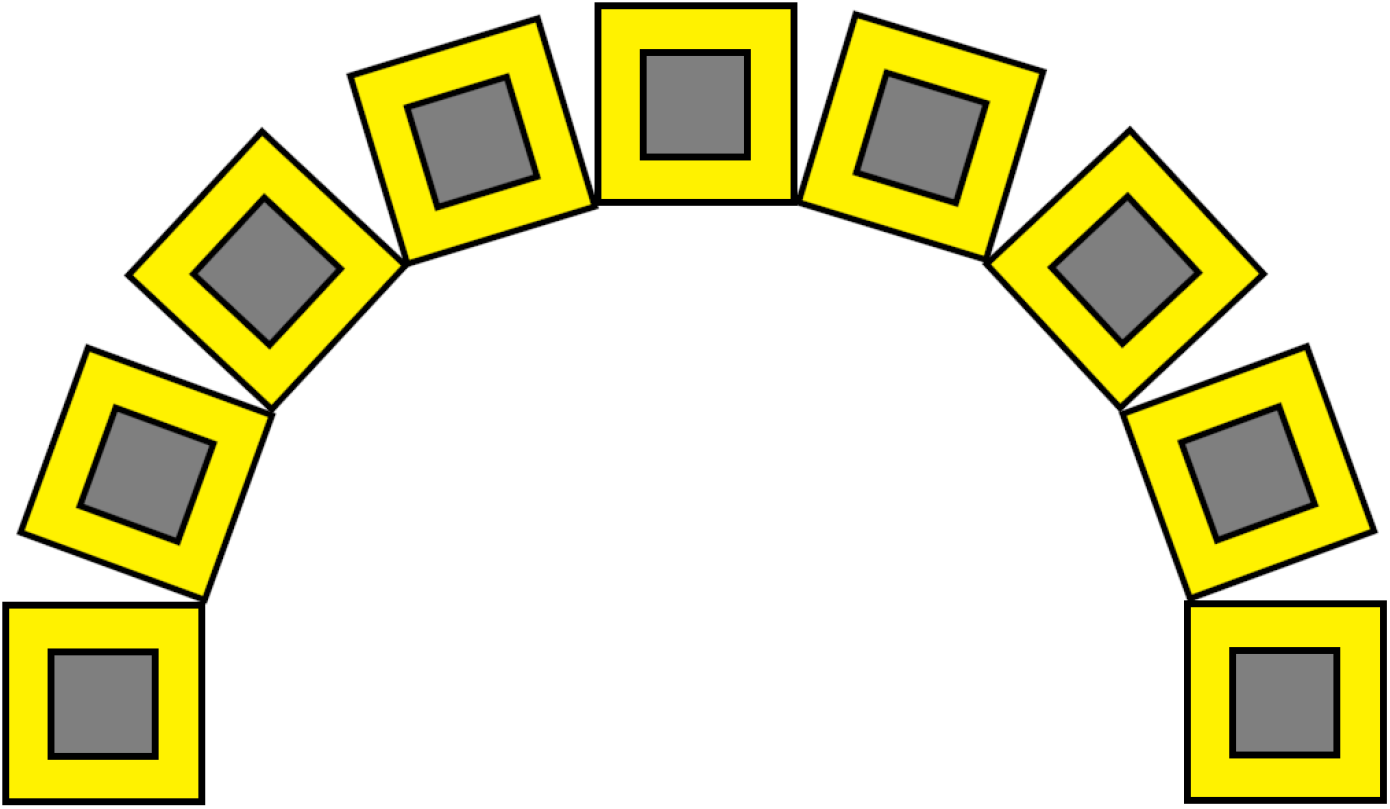
Top-down view of the setting. The apparatuses are represented as grey squares, and the tables that support them as yellow squares. In this diagram, the entrance to the enclosure would be located below the bottom of the figure.

The boxes were made of Plexiglas bolted together with corner brackets to prevent the animals from damaging them. The Plexiglas used for the wolf boxes had a thickness of 1cm to make the boxes more durable and thus reduce the possibility of the animals breaking them. Several holes drilled through the walls of the boxes allowed the animals to smell the reward from the outside, to entice them to interact with them. The boxes were semi-transparent (with thin, diagonal dark lines painted across all their walls) to account for potential differences in their experience with transparent objects. A schematic of all the apparatuses can be seen in Figure 2, and Supplementary Video S1.

**Figure 2.**
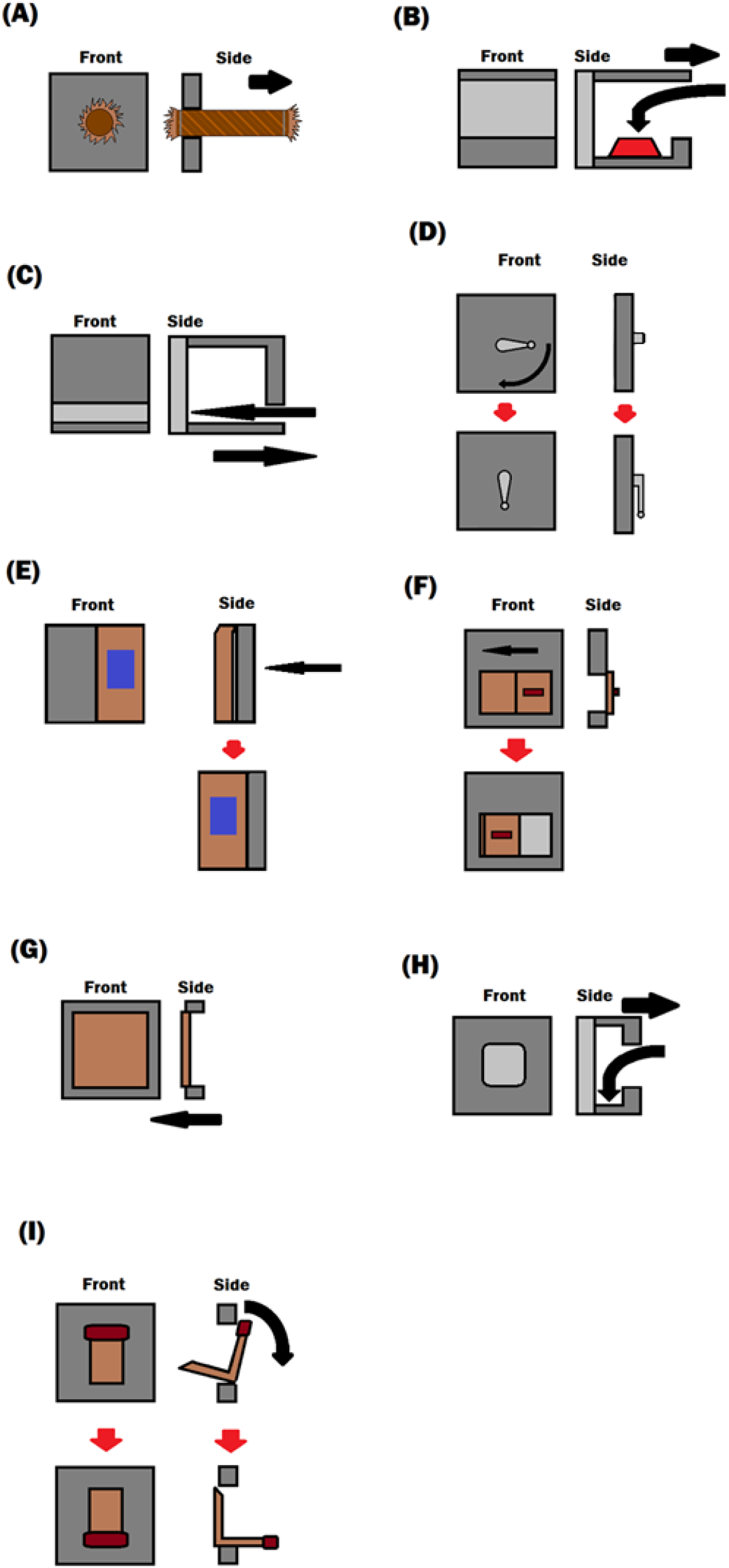
Front and side views of the apparatuses used in this experiment (other parts of the box were excluded if there were not relevant). The black arrows indicate the direction of the movement the animals had to make to obtain the reward, while the red arrows (when present) indicate the changes between the “unsolved” and “solved” configurations of a given apparatus. **(A)** Rope. **(B)** Bowl. **(C)** Slit. **(D)** Door handle. **(E)** Push. **(F)** Slide door. **(G)** Breakable. **(H)** Double tray. **(I)** Flip box.

#### Rope (Figure 2, (A))

The rope was tied to the door of the box. By pulling the rope, the subject was able to open the box.

#### Bowl (Figure 2, (B))

The box had a wide opening in the front wall, with a small plexiglass strip covering the bottom. Inside the box, an upside-down bowl (a small lunch box for the dogs and a metallic sieve for the wolves, both of which with holes to allow for the smell to pass through) covered the reward. The subjects had to either flip the bowl with their muzzle or paws, or to simply take out the bowl altogether to uncover the reward. The bottom strip of Plexiglas on the front of the box prevented the subjects from simply sliding the bowl all the way out of the box without the need to flip it or remove it in any manner.

#### Slit (Figure 2, (C))

The box had a narrow slit on the front wall, with no other features. To reach the reward, the subjects had to stick one of their paws inside and pull towards themselves. To account for the different sizes of the animals, the reward was placed slightly closer to the entry for the dogs.

#### Door handle (Figure 2, (D))

A door handle measuring 12 cm was placed on the front wall of the box, with the lever parallel to the to that wall itself. Rotating the handle (that is, putting it pointed towards the bottom of the box) opened the front wall of the box (fixed to one of the sides with hinges).

#### Push (Figure 2, (E))

Half of the front of the box was attached to the side with hinges in such a manner that it could be pushed inside by gently pressing it, thus making the reward in the box (placed near one of the corners to prevent the subjects from sweeping it out of their reach) accessible to the animal. A blue rectangle was painted on the openable side to make it easily distinguishable.

#### Slide door (Figure 2, (F))

The front wall of the box was provided with a hole, covered by a sliding door on metal rails. This door had a small handle protruding approximately 2 cm that the subjects could either grab with their mouth or push with their paws. By moving the door to the side, the opening to the box (and with it, the reward) became accessible to the subject to reach with their paws or muzzle.

#### Breakable (Figure 2, (G))

The front of the box was covered by a breakable material (newspapers for wolves and paper tissues for dogs, attending to structural considerations due to the different sizes of the boxes). By breaking the cover, the subject had direct access to the reward.

#### Double tray (Figure 2, (H))

The front side of the box was closed with two panes of Plexiglas, the inner of which was connected to the rest of the box through a hinge. The outer pane had a hole in the center (large enough to allow the animals to stick their paws inside) while the inner one was completely solid. By putting their paws in between the two panes and pulling towards themselves, the subjects were able to pull the whole structure blocking the opening of the box to the side to access the reward.

#### Flip box (Figure 2, (I))

A smaller box shaped like a rectangular triangle prism with the top side open was placed inside the box, with a handle protruding towards the front of the puzzle box. By pulling down the handle, the open side of the flip box passed through the front of the puzzle box, becoming accessible to the subject.

### Procedure

All sessions took place within the same outdoor enclosure at the Wolf Science Center, measuring 620 m^2^. Regardless of the type of session (habituation or testing), each testing day started with 5 minutes of free exploration of the enclosure to prevent the animals from getting distracted by the smell of other animals that had recently stayed in that enclosure. After this, the subjects were taken out of the enclosure through the shifting system^1^ and the session was prepared.

#### Habituation

To increase the likelihood of all animals (wolves and dogs) approaching the boxes, a habituation session took place. A single, featureless box without a front side was placed inside the enclosure while the subject waited in the shifting system. A trainer that had established a relationship with the subject then walked to the box and put several food rewards inside it. After this, the subject was released into the enclosure and allowed to reach the food inside the box, either by placing their muzzle inside of the apparatus or fishing for the reward with their paws. The apparatus was re-baited by the trainer several times during this session, until the subject reliably retrieved the reward on their own.

If an animal consistently failed to retrieve the reward and showed avoidance towards the apparatus during this session, the habituation session was repeated on a different day, up to a maximum of two times. Only two subjects (two wolves) required more than one habituation session before moving on to the test phase (see **Table 1** for more details). In any case, the first testing session took place immediately after a subject had successfully completed the habituation session.

#### Test

While the subject was inside the shifting system, the 9 test apparatuses were placed inside the enclosure, describing a semi-circle with the open side facing the entrance through which the animal would enter the enclosure (see **Figure 1**), so that they were all equidistant to the subject upon reaching the grid of apparatuses — defined as the area described by the semi-circle of apparatuses. As the apparatuses for wolves and dogs had different sizes (see above), the size of the semicircle differed between the species (2m for the dogs, 4 meters for the wolves), as well as the distance between the center of the semicircle and the entrance of the enclosure (4.5 meters for the dogs, 5.5 meters for the wolves. All apparatuses were baited with a single food reward (a small piece of meat).

After all the apparatuses were ready, the subject was released into the enclosure (this time, without the trainer being present) and allowed to freely interact with each of the apparatuses for a maximum of 9 minutes (with 1 minute allotted to solve each of the apparatuses), at which point, the session ended and the animal was called out of the test enclosure. A session would also end whenever a subject solved all 9 apparatuses in less than 9 minutes. Each subject was exposed to 10 of such test sessions (except for one subject that refused to enter the test enclosure in one of the sessions, and another one in which the experiment was interrupted by heavy rain; see **Table 1** for more details). All testing sessions took place on different days, usually in a regimen of 2-3 sessions per week for each animal (aside from some cases with the pet dogs, as their participation in the study depended on their owner’s availability).

## Analyses

All sessions were filmed with two camcorders to cover the whole breadth of the array of apparatuses. The resulting videos were later coded with BORIS® (Friard & Gamba, 2016, http://www.boris.unito.it/) by someone blind to the hypotheses. A fraction of the videos, pertaining to 56 sessions (randomized between all three populations) were also coded by one of the experimenters to assess inter-rater reliability. We used the intra-class correlation statistic (for continuous variables) and Cohen’s kappa (for binary variables), both of which were calculated with the “irr” package (Gamer & Lemon, 2019), within R 4.0.2 (R Core Team, 2013). Inter-rater agreement was above 0.9 for all variables used for data analyses.

To compare our results with previous experiments, we originally planned to design our statistical analyses after Rössler & Mioduszewska et al. (2020), performing a Principal Component Analysis (PCA) on 6 variables related to the animals’ performance. However, we found some issues both in the statistical design of the study and in the distribution of the variables that were used for the PCA, prompting us to remove half of those variables (see Supplementary Materials for more information). The remaining variables were mostly correlated to each other, thus making the use of a PCA no longer appropriate.

As such, we used Linear Mixed Models (LMM) to analyze the differences in performance across sessions and between all three populations both in terms of proportion of time interacting with the apparatuses (Models 1.1 and 1.2 in **Table 2**) and the latency to reach the grid of apparatuses (Models 2.1 and 2.2 in **Table 2**). Proportion of time interacting with the apparatuses was logit-transformed (as per Warton & Hui, 2011), with zero values being replaced by the lowest non-zero value registered for that measurement divided by two before the transformation. Latency to approach the grid of apparatuses was log-transformed, with a value of 540 seconds (9 minutes) being used for those sessions in which the subject did not approach the apparatuses.

**Table 2.**
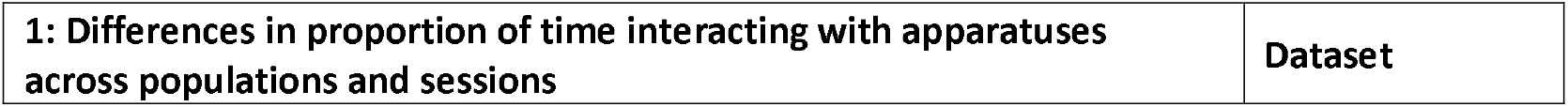
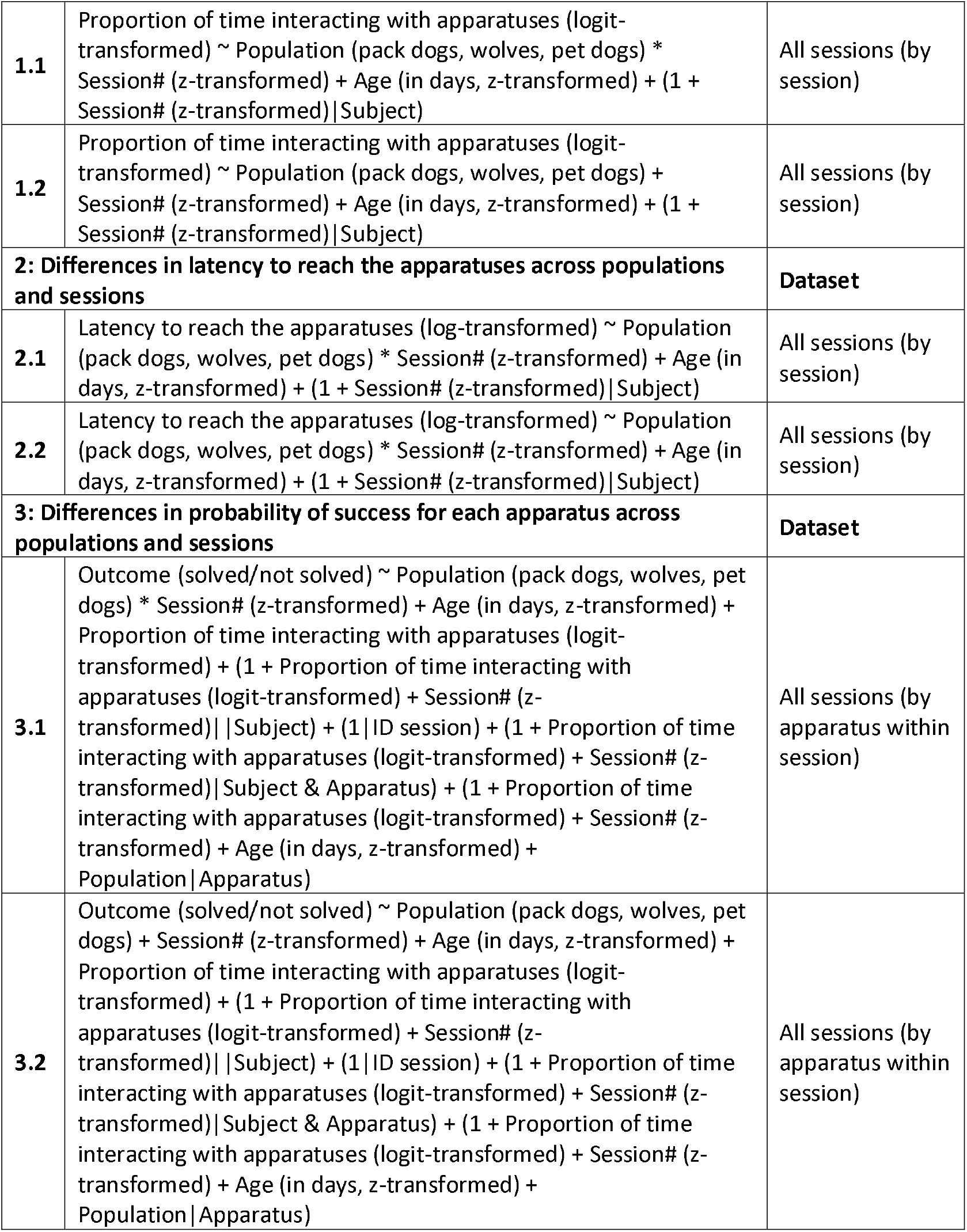
Models used in the experiment.

Afterwards, similar to the analyses done by Rössler, Mioduszewska and colleagues, we calculated the probability to solve each of the boxes across sessions and between all three populations through a Generalized Linear Mixed Models (GLMM) with binomial error structure and logit link function. The original study included the principal components of the PCA as fixed effects, to test for how different performance-related factors influenced probability of solving the boxes. To mirror this, we included the proportion of time interacting with the apparatuses as a fixed effect in our model (Models 3.1 and 3.2 on **Table 2**). Since the latency to reach the apparatuses was correlated with the proportion of time interacting with them (Pearson’s correlation: t(322) = −16.51, R^2^ = 0.458, p < 0.001), we decided to exclude this variable from the analyses.

Since exploration patterns are known to change with age in wolves and dogs (Jean-Joseph et al., 2025; Moretti, 2015), we included the age of the subjects in days as a continuous control variable in all models.

All models were calculated by using the lme4 package (V.0.6.7, Bates et al., 2015). The degrees of freedom for the linear mixed models were calculated by using Satterthwaite’s method through the lmerTest (V.0.8.2, Kuznetsova et al., 2017) package.

Random slopes were added whenever there was enough variation in those variables for each of the levels of the random intercept. The model stability was analyzed through a custom function devised by Roger Mundry and modified by Remco Folkertsma. The collinearity was tested through the “vif” function of the “car” (Fox and Weisberg, 2018) package. We compared all models with a null model lacking the predictor variables of the full model but otherwise containing the same random effects and control variables. In all models, we z-transformed all continuous predictors to obtain more easily interpretable model estimates and prevent model convergence issues. Then, we used the Anova function to compare the full-null models with the Chi-squared test. A detailed description of each of the models used can be found in **Table 2**.

## Ethical statement

All procedures were approved by the “Ethics and Animal Welfare Committee at the University of Veterinary Medicine Vienna” with the ETK number “ETK-164/11/2020”. All animals tested were either wolves and dogs raised and living in captivity or pet dogs (therefore, no wild animals were tested). Participation in the experiment was voluntary for all animals. None of the subjects (neither the captive wolves and dogs nor the pet dogs) were food deprived, and they had permanent access to water during testing sessions. Regarding the pet dogs, the owners’ consent was given in advance, and said owners were also informed beforehand about the procedure of the study. At any time, the animals had the possibility to retreat into the enclosure in case they were uncomfortable with the experiments. We followed ASAB (Guidelines for the Ethical Treatment of Nonhuman Animals in Behavioural Research and Teaching, 2023) guidelines for the ethical treatment of animals in behavioral research and teaching.

## Results

Animals showed a great degree of variability in their frequency of interaction with the apparatuses. One wolf (*Taima*) never entered the apparatus-grid, while 4 other wolves (*Kenai, Nanuk, Tekoa*, and *Una*) and 3 pack dogs (*Enzi, Layla*, and *Panya*) rarely interacted with the apparatuses and never solved any of them. Conversely, all pet dogs solved at least 1 apparatus (albeit one of them — *Pepeo*— did so only once, and showed little interest in the task throughout the sessions).

Several animals managed to solve each of the apparatuses in at least 1 session, totaling 4 wolves (*Amarok, Etu, Tala*, and *Yukon*), 4 pack dogs (*Hiari*, Imara, Moya, and *Inaya*), and 6 pet dogs (*Asali, Bora, Hakima, Zazu, Zhenga*, and *Freya*). Indeed, pet dogs had the highest proportion of animals solving all boxes: out of the 3 pet dogs that did not solve all of them, 2 of them (*Kilio* and *Koda*) solved 8 apparatuses in at least one session. A detailed account of the number of apparatuses solved by each animal in each session can be found in **Supplementary Table S2**, and the number of sessions in which each subject solved each one of the boxes can be found in **Supplementary Table S3**.

All 9 apparatuses were solved by at least one animal in all three populations, albeit some of them were solved more than others. Although there was variation between populations, some of the apparatuses —such as “slit” and “breakable”— were solved by most of the animals, while some others (such as “double tray” and “door handle”) were rarely solved, both in terms of raw number of sessions and subjects that managed to solve them. **Supplementary Table S4** provides a full account of the likelihood of solving each apparatus per population, as calculated by model 3.1 (**Table 2**).

### Interaction time with the apparatuses

The model analyzing the proportion of time the animals spent interacting with the apparatuses was not significantly different to the null model (Model 1.1 on **Table 2**; full-null comparison: χ^2^ = 4.511, p = 0.478). This remained the case for the reduced model that lacked the interaction between population and session (Model 1.2 on **Table 2**; full-null comparison: χ^2^ = 4.495, p = 0.213). This indicates that there were no significant differences between the populations in the amount of time they interacted with the boxes nor how this variable changed throughout the experiment.

Despite this, the individuals with the lowest average interaction times were consistently wolves and pack dogs, with only one pet dog (*Pepeo*) among the bottom 50% of individuals in terms of interaction time (see **Supplementary Table S5** for more details).

### Latency to reach the grid of apparatuses

The model with the latency to approach the grid of apparatuses as the response variable was significantly different from the null (Model 2.1 on **Table 2**; full-null comparison: χ2 = 186.634, p < 0.001). The model showed that wolves needed more time than pack dogs to approach the apparatuses (estimate ± SE = 1.287 ± 0.519, t(34.665) = 2.473, p = 0.018). As the interaction between population and session proved to be non-significant however, we ran the reduced model without this interaction. This model was also different from the null (Model 2.2 on **Table 2**; full-null comparison: χ2 = 185.865, p < 0.001), and it showed once again that wolves needed more time than pack dogs to approach the apparatuses (estimate ± SE = 1.447 ± 0.485, t(32.839) = 2.984, p = 0.005).

**Figure 3.**
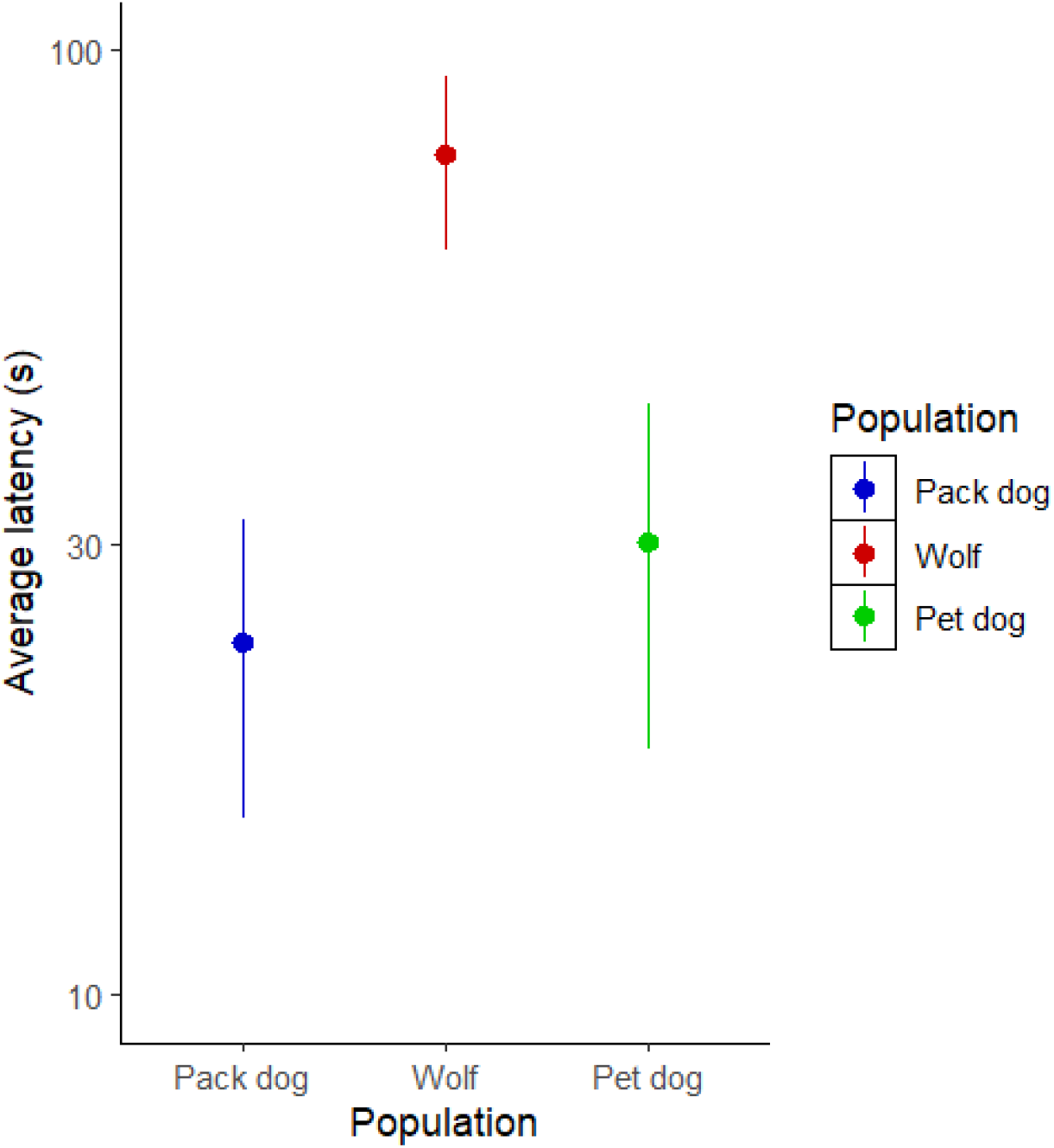
Average latency to reach the grid of apparatuses per population, with length of the whiskers representing the standard error of the measurements. The y-axis has been log-transformed for clarity.

### Probability of solving the apparatuses

The model we used to investigate the factors affecting the probability to solve each apparatus was significantly different from the null model (full-null comparison: χ^2^ = 269.847, p < 0.001). The proportion of time interacting with the boxes was significant, with more interaction leading to a higher probability of success (estimate ± SE = 1.837 ± 0.294, z = 6.252, p < 0.001).The effect of the session was significant as well (estimate ± SE = 1.357 ± 0.385, z = 3.524, p < 0.001), but not the interaction between session and population (effect of session in pack dogs vs wolves: estimate ± SE = −0.480 ± 0.319, z = −1.508, p = 0.132; effect of session in pack dogs vs pet dogs: estimate ± SE = 0.386 ± 0.419, z = 0.922, p = 0.356), which suggests that the probability of solving an apparatus increased in each session in a similar manner for all three populations. To further investigate this, we fitted a reduced model with the same variables as the full one except for the interaction between population and session. Once again, this model was significantly different from the null model (reduced-null comparison: χ^2^ = 264.720, p < 0.001), and the interaction time with the apparatuses had a significant effect (estimate ± SE = 1.726 ± 0.266, z = 6.499, p < 0.001). As expected, the effect of the session was still present (estimate ± SE = 1.489 ± 0.219, z = 6.801, p < 0.001), but this time we also found an effect of population, with pet dogs being more successful at solving the apparatuses than the pack dogs (estimate ± SE = 5.233 ± 2.031, z = 2.577, p = 0.010). The age of the animals also proved to have a significant effect, with older individuals having a lower probability of success, even when controlling for interaction time (estimate ± SE = −1.980 ± 0.866, z = −2.285, p = 0.022).

Probability of success solving each given apparatus varied depending on the identity of the apparatus and the species, although most of the apparatuses were solved at a similar relative rate between all three populations. A full report of each apparatus’ probability of being solved by each population, as well as its difficulty rank, can be found in **Supplementary Table S4**.

**Figure 4.**
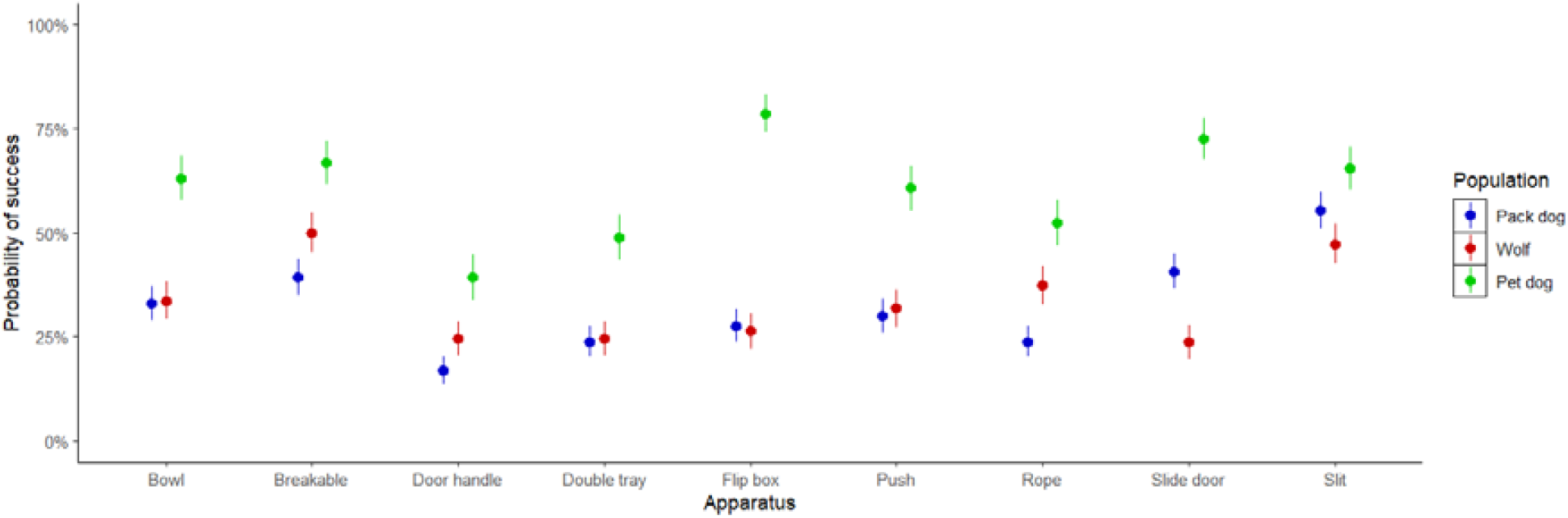
Average probability of success on each apparatus for all three populations. Whiskers represent the standard error of this measurement.

**Figure 5.**
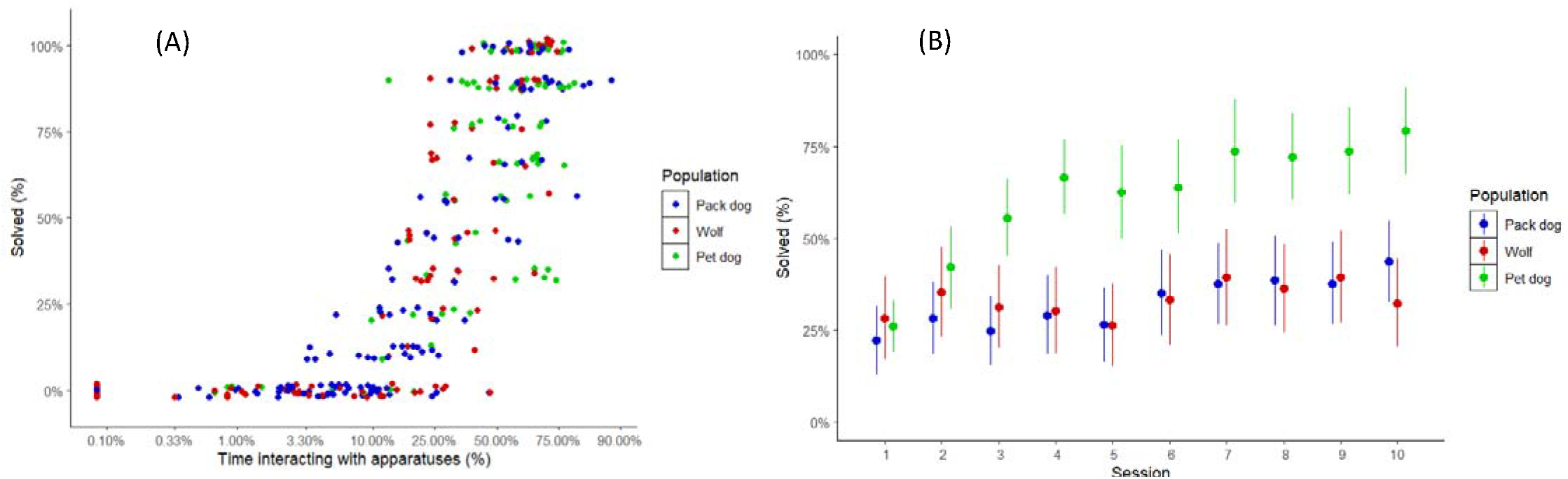
Percentage of boxes solved per session as a factor of two variables: (A) Time interacting with boxes (as percentage of the total session time) (B) Session number. Individuals points in plot (A) represent total interaction time in a given session (for clarity, the x axis has been scaled to the logit function). Values for plot (B) are the average of the number of apparatuses solved in all sessions of the same number across all individuals of each population (the value used for the whiskers in the plot is the standard error of that measurement).

## Discussion

In this study, we were set to explore the differences in the way wolves and dogs engage with innovative problem-solving tasks, as well as how these different approaches relate to problem solving success. We found no differences between pack dogs and wolves and pack dogs and pet dogs in the amount of time they spent interacting with the apparatuses, but we found that wolves approached the apparatuses slower than pack dogs. Despite this lack of differences in interaction, pet dogs were more successful at solving the boxes than pack dogs. Indeed, all but 1 of the pet dogs solved at least 8 out of the 9 apparatuses, and all but 3 of the pet dogs solved all 9.

Overall, pet dogs’ higher rate of success matches our predictions, as we hypothesized that pet dogs’ previous positive experiences with human-created objects would prove advantageous for them (as discussed by Lazzaroni et al., 2019). Pet dogs’ performance could have come as the result of prior experience with situations in their daily life, for example, prior usage of “cognitive toys” —which anecdotally, some of the animals did have according to their owners’ report. Having been able to get food from similar contraptions or/and encouragement by their owners may have prompted them to make a greater effort in getting the reward. Alternatively, previous experience with physical principles of other human-created constructs, and having a greater positive experience with object-directed play may have also had a role. Both the prior positive experiences and any generalization of their physical attributes could have been enhanced by the fact that all pet dogs we tested belonged to trainers working at the Wolf Science Center, which means they likely engaged in object-directed play and similar activities with their pets to a higher rate than the average pet owner. Thus, results of the present study ought to be taken with caution in what pertains to the pet dogs, as their prior experiences may be far richer than the average pet dogs’.

Rössler, Mioduszewska, and colleagues (2020) observed in their study that some of the wild Goffin cockatoos showed little interest in interacting with the apparatuses, regardless of the fact that they found no differences in problem solving success among motivated individuals. They hypothesized that the captive population may have been more motivated due to the expectation of receiving a reward because of their previous experience participating in other studies in the lab. In the current study all three populations should have had a similar expectation. However, the animals that interacted the least with the apparatuses in our study were almost exclusively wolves and pack dogs. This could imply that, rather than previous experience with being tested, the extent to which they were habituated to human-created constructs prompted both cockatoos and canids to interact with the apparatuses. Previous history of reinforcement with human-created constructs could, then, serve as an initial driver to explore similar novel objects, which would eventually lead to the discovery of novel opportunities.

Our results seem to fall in line with an impossible task study performed by Lazzaroni et al. (2019). In it, three populations of dogs were tested: two of them equivalent to the ones used in the current study, and a third one composed of free-ranging dogs, who had prior limited contact with human-created constructs. In a similar vein to both our study and the one done by Rössler, Mioduszewska, and colleagues (2020), the population with the least prior positive experiences with human-created objects was the one that showed the least interest in the task. Similar to our study as well, Lazzaroni and colleagues found no overall differences in terms of interaction time between pack dogs and pet dogs. However, contrasting our study, pet dogs did not show a greater motivation to participate in the task, with roughly equivalent numbers of individuals that failed to interact with the apparatus between pet and pack dogs. Therefore, although animals with the most positive prior experiences with human-created objects seem in general more prone to interact with novel objects, the specifics of this apparent trend may ultimately come down to individual experiences and the nature of the objects at hand.

Somewhat matching our predictions, wolves showed a higher latency to approach the apparatuses when compared with pack dogs, potentially due to neophobia. Interestingly, differences in latency to approach did not decrease across sessions. They also did not translate into differences between the species in terms of persistence of exploration and, most importantly, problem-solving success. This lack of differences in innovative problem-solving skills between the species could respond to one of four possibilities. The first scenario suggests that distinct evolutionary pressures facilitated innovation in wolves and dogs. Dogs’ evolutionary environment was likely highly diverse but generally abundant in resources, while wolves’ was much harsher, but also less diverse. Thus, both pressures may have potentially reached the same point. A second, more parsimonious scenario would point towards no differences arising due to the relatively recent split between both species. Indeed, wolves and dogs only started to diverge around 30 000 years ago (Cooper et al., 2015), which may have not been time enough for differences in innovative problem solving to arise. A third scenario suggests that both species’ behavioral repertoire is too limited to see any advantages of innovative problem solving, and thus, no selective pressure acted on that trait. As opposed to other species known for their innovative problem solving such as apes and some birds, dogs and wolves have a restricted repertoire in what pertains to the manipulation of objects, mostly circumscribed to biting and scratching. Such behaviors may have not been useful enough to exploit novel opportunities in these species’ evolutionary past, (as discussed by Tebbich et al., 2016). Finally, a fourth scenario suggests that the effect of experience was strong enough to mask any differences in behavior caused by each species’ evolutionary history. Although pet dogs were the population with the most experience with human-created objects, the wolves and pack dogs were also familiar with these objects due to their life in captivity and participation in previous experiments. It could, then, be the case that naïve wolves and dogs would show differences in their innovative problem-solving behavior shaped by their evolutionary history.

The relative ease with which each of the apparatuses was solved remained mostly the same among all three populations, with the most straightforward ones —such as breakable and slit— being among the easiest to solve for all three populations, while the ones with most complicated mechanisms —such as double tray and door handle— coming in the last few slots in all three populations. Four of the apparatuses, however, showed higher differences in solvability between the populations: slide door, door handle, push, and flip box. In the case of the slide door and door handle apparatuses, the observed differences in performance may have come as a result of structural differences between the apparatuses of the wolves and the dogs. In case of the dogs, several animals managed to move the sliding door by merely pawing at the front of the apparatus. Conversely, wolves needed to apply force laterally (either by pushing or biting the handle) in order to open the door. Thus, it should not be surprising that wolves solved it to a lesser extent than the two other populations. The opposite happened with the door handle mechanism, which was more structurally stable in the case of the wolf boxes, and thus, responded better to the animals’ attempts to manipulate it. In case of the rope apparatus, the likely explanation for the differences in performance between the populations is that it may have come as a result of different prior experiences. Indeed, all wolves had previous experience pulling from ropes from previous experiments (e.g.: Marshall-Pescini, 2018; Range et al., 2019a; Range et al., 2019b), but only a limited subset of both pet and pack dogs had done so in the past. This seems to match our results, as wolves solved it to a greater degree than either population of dogs did. The remaining apparatus with differences between the species (flip box) seems harder to explain, but seeing as it was the one that was solved the most by pet dogs, whose life experiences should be the most unique among all three populations, this may also have to do with prior experiences with similar contraptions.

In conclusion, we did find an effect of prior experience with human-created constructs in dogs’ innovative problem-solving abilities, but no apparent effect of domestication given packs dogs’ similar performance to that of wolves. Animals’ prior experience did not only impact their performance, but also whether they were motivated in interacting with the puzzle boxes in the first place.

## Author contributions

FR, ST and SMP conceptualized the experiment and secured funding; DRB developed the methodology and crafted the apparatuses for the dogs; FVS and DRB performed the data collection; DRB curated and analyzed the data; FVS coded a fraction of the videos; DRB wrote the first draft of the paper; FR, ST and SMP supervised the entirety of the experiment; all authors edited and reviewed the manuscript.

## Acknowledgements

We thank Kathryn Roddie, Lou Gonnet-dit-Revel and the trainers at the Wolf Science Center Core facility for contributing to the data collection process, Juliette Gratalon for coding of the videos, and Erik Hollerweger for crafting the apparatuses for the wolves and the tables for both dogs’ and wolves’ apparatuses. We also thank Theresa Rössler and Remco Folkertsma for their help with the statistical analyses.

## Footnotes

1 All enclosures are surrounded by a fenced corridor where the animals can be placed in order to perform maintenance activities or move the animals to adjacent enclosures.

