## Supplementary text, tables, and figures for "Figuring things out: Innovation arena in wolves and dogs"

### Supplementary materials

**Supplementary Table S1:** Details about the subjects participating in this experiment.

| Subject | Species / Population | Subspecies / Breed | Relatives | Age (months) | Sex |
| --- | --- | --- | --- | --- | --- |
| Amarok | Wolf | Timber wolf | Tala | 112 | Male |
| Chitto |  | Timber wolf | Una | 111 | Female |
| Etu |  | Timber wolf |  | 64 | Male |
| Geronimo |  | Timber wolf | Yukon | 146 | Female |
| Kenai |  | Hudson Bay wolf |  | 136 | Male |
| Maikan |  | Timber + Eurasian wolf | Tekoa, Taima | 64 | Male |
| Nanuk |  | Timber wolf |  | 147 | Male |
| Taima |  | Timber + Eurasian wolf | Tekoa, Maikan | 64 | Female |
| Tala |  | Timber wolf | Amarok | 111 | Male |
| Tekoa |  | Timber + Eurasian wolf | Taima, Maikan | 64 | Male |
| Una |  | Timber wolf | Chitto | 112 | Male |
| Yukon |  | Timber wolf | Geronimo | 146 | Male |
| Enzi | Pack dog | Mongrel | Bora, Layla, Panya, Zazu, Pepeo | 84 | Male |
| Hiari |  | Mongrel | Imara | 86 | Male |
| Imara |  | Mongrel | Hiari | 86 | Female |
| Inaya |  | Mongrel | Kiano | 11 | Female |
| Kiano |  | Mongrel | Inaya | 11 | Male |
| Kovu |  | Mongrel | Youma | 14 | Male |
| Layla |  | Mongrel | Bora, Enzi, Panya, Zazu, Pepeo | 118 | Female |
| Moya |  | Mongrel | Sarabi, Taio | 9 | Female |
| Taio |  | Mongrel | Moya, Sarabi | 11 | Male |
| Sarabi |  | Mongrel | Moya, Taio | 9 | Female |
| Panya |  | Mongrel | Bora, Layla, Enzi, Zazu, Pepeo | 86 | Female |
| Youma |  | Mongrel | Kovu | 14 | Female |
| Zuri |  | Mongrel |  | 118 | Female |
| Asali | Pet dog | Mongrel† |  | 127 | Male |
| Bora |  | Mongrel | Layla, Enzi, Panya, Pepeo, Zazu | 118 | Female |
| Freya |  | Irish Terrier |  | 72 | Female |
| Hakima |  | Mongrel† |  | 127 | Male* |
| Kilio |  | Mongrel† |  | 136 | Male* |
| Koda |  | Mixed-breed |  | 121 | Male |
| Pepeo |  | Mongrel | Bora, Layla, Enzi, Panya, Pepeo | 85 | Male |
| Zazu |  | Mongrel | Bora, Layla, Enzi, Panya, Pepeo | 84 | Male* |
| Zhenga |  | Australian shepherd |  | 48 | Female |

* Castrated.
† Related to other WSC dogs not participating in this experiment.

**Supplementary Table S2:** Number of sessions in which each of the apparatuses was solved per animal. Greener colors represent a higher number, while redder ones represent a lower number.

|  |  |  | **Bowl** | **Breakable** | **Door handle** | **Double tray** | **Flip box** | **Push** | **Rope** | **Slide door** | **Slit** |
| --- | --- | --- | --- | --- | --- | --- | --- | --- | --- | --- | --- |
| **Amarok** | **Wolf** | | 6 | 10 | 10 | 10 | 9 | 10 | 10 | 10 | 10 |
| **Chitto** | **Wolf** | | 3 | 10 | 1 | 2 | 2 | 1 | 2 | 0 | 7 |
| **Etu** | **Wolf** | | 10 | 10 | 10 | 9 | 8 | 9 | 10 | 10 | 10 |
| **Geronimo** | **Wolf** | | 6 | 10 | 0 | 0 | 0 | 2 | 7 | 0 | 6 |
| **Kenai** | **Wolf** | | 0 | 0 | 0 | 0 | 0 | 0 | 0 | 0 | 0 |
| **Maikan** | **Wolf** | | 0 | 0 | 0 | 0 | 0 | 0 | 0 | 0 | 0 |
| **Nanuk** | **Wolf** | | 0 | 0 | 0 | 0 | 0 | 0 | 0 | 0 | 0 |
| **Taima** | **Wolf** | | 0 | 0 | 0 | 0 | 0 | 0 | 0 | 0 | 0 |
| **Tala** | **Wolf** | | 10 | 10 | 5 | 4 | 8 | 9 | 10 | 2 | 10 |
| **Tekoa** | **Wolf** | | 0 | 0 | 0 | 0 | 0 | 0 | 0 | 0 | 0 |
| **Una** | **Wolf** | | 0 | 0 | 0 | 0 | 0 | 0 | 0 | 0 | 0 |
| **Yukon** | **Wolf** | | 2 | 5 | 1 | 2 | 2 | 4 | 2 | 4 | 9 |
| **Enzi** | **Dog** | **Pack** | 0 | 0 | 0 | 0 | 0 | 0 | 0 | 0 | 0 |
| **Hiari** | **Dog** | **Pack** | 10 | 10 | 9 | 9 | 10 | 8 | 10 | 10 | 10 |
| **Imara** | **Dog** | **Pack** | 8 | 10 | 3 | 9 | 10 | 8 | 8 | 10 | 10 |
| **Layla** | **Dog** | **Pack** | 0 | 0 | 0 | 0 | 0 | 0 | 0 | 0 | 0 |
| **Panya** | **Dog** | **Pack** | 0 | 0 | 0 | 0 | 0 | 0 | 0 | 0 | 0 |
| **Zuri** | **Dog** | **Pack** | 0 | 0 | 2 | 1 | 0 | 2 | 0 | 2 | 4 |
| **Moya** | **Dog** | **Pack** | 10 | 9 | 4 | 6 | 8 | 8 | 9 | 10 | 10 |
| **Taio** | **Dog** | **Pack** | 0 | 7 | 0 | 0 | 1 | 2 | 0 | 2 | 7 |
| **Sarabi** | **Dog** | **Pack** | 6 | 8 | 0 | 0 | 0 | 3 | 0 | 10 | 9 |
| **Kiano** | **Dog** | **Pack** | 0 | 0 | 0 | 0 | 0 | 0 | 0 | 0 | 5 |
| **Inaya** | **Dog** | **Pack** | 9 | 7 | 4 | 6 | 7 | 7 | 4 | 9 | 10 |
| **Kovu** | **Dog** | **Pack** | 0 | 0 | 0 | 0 | 0 | 1 | 0 | 0 | 4 |
| **Youma** | **Dog** | **Pack** | 0 | 1 | 0 | 0 | 0 | 0 | 0 | 0 | 4 |
| **Asali** | **Dog** | **Pet** | 3 | 8 | 5 | 9 | 8 | 6 | 3 | 8 | 2 |
| **Bora** | **Dog** | **Pet** | 8 | 7 | 5 | 5 | 9 | 8 | 9 | 8 | 6 |
| **Hakima** | **Dog** | **Pet** | 4 | 7 | 2 | 4 | 10 | 6 | 4 | 7 | 10 |
| **Kilio** | **Dog** | **Pet** | 9 | 4 | 0 | 1 | 7 | 4 | 2 | 9 | 8 |
| **Pepeo** | **Dog** | **Pet** | 0 | 0 | 0 | 0 | 0 | 0 | 1 | 0 | 0 |
| **Zazu** | **Dog** | **Pet** | 9 | 9 | 8 | 9 | 10 | 7 | 10 | 9 | 8 |
| **Zhenga** | **Dog** | **Pet** | 8 | 9 | 8 | 6 | 10 | 8 | 6 | 8 | 9 |
| **Koda** | **Dog** | **Pet** | 4 | 4 | 1 | 0 | 4 | 4 | 1 | 4 | 4 |
| **Freya** | **Dog** | **Pet** | 8 | 8 | 4 | 7 | 8 | 8 | 8 | 8 | 8 |

**Supplementary Table S3:** Number of apparatuses solved per animal and session. Greener colors represent a higher number, while redder ones represent a lower number.

|  |  |  | **1** | **2** | **3** | **4** | **5** | **6** | **7** | **8** | **9** | **10** |
| --- | --- | --- | --- | --- | --- | --- | --- | --- | --- | --- | --- | --- |
| **Amarok** | **Wolf** | | 9 | 9 | 7 | 8 | 8 | 8 | 9 | 9 | 9 | 9 |
| **Chitto** | **Wolf** | | 3 | 3 | 2 | 2 | 1 | 1 | 4 | 3 | 6 | 3 |
| **Etu** | **Wolf** | | 8 | 8 | 9 | 9 | 8 | 9 | 9 | 9 | 9 | 8 |
| **Geronimo** | **Wolf** | | 1 | 3 | 3 | 2 | 3 | 4 | 4 | 4 | 3 | 4 |
| **Kenai** | **Wolf** | | 0 | 0 | 0 | 0 | 0 | 0 | 0 | 0 | 0 | 0 |
| **Maikan** | **Wolf** | |  |  |  |  |  |  |  |  |  |  |
| **Nanuk** | **Wolf** | | 0 | 0 | 0 | 0 | 0 | 0 | 0 | 0 | 0 | 0 |
| **Taima** | **Wolf** | | 0 | 0 | 0 | 0 | 0 | 0 | 0 | 0 | 0 | 0 |
| **Tala** | **Wolf** | | 5 | 8 | 7 | 7 | 6 | 8 | 9 | 5 | 6 | 7 |
| **Tekoa** | **Wolf** | | 0 | 0 | 0 | 0 | 0 | 0 | 0 | 0 | 0 | 0 |
| **Una** | **Wolf** | | 0 | 0 | 0 | 0 | 0 | 0 | 0 | 0 | 0 | 0 |
| **Yukon** | **Wolf** | | 2 | 4 | 3 | 2 | 0 | 3 | 4 | 6 | 6 | 1 |
| **Enzi** | **Dog** | **Pack** | 0 | 0 | 0 | 0 | 0 | 0 | 0 | 0 | 0 | 0 |
| **Hiari** | **Dog** | **Pack** | 8 | 9 | 8 | 9 | 9 | 9 | 8 | 9 | 8 | 9 |
| **Imara** | **Dog** | **Pack** | 8 | 7 | 7 | 6 | 7 | 8 | 8 | 9 | 8 | 8 |
| **Layla** | **Dog** | **Pack** | 0 | 0 | 0 | 0 | 0 | 0 | 0 | 0 | 0 | 0 |
| **Panya** | **Dog** | **Pack** | 0 | 0 | 0 | 0 | 0 | 0 | 0 | 0 | 0 | 0 |
| **Zuri** | **Dog** | **Pack** | 0 | 3 | 1 | 0 | 0 | 1 | 3 | 0 | 1 | 2 |
| **Moya** | **Dog** | **Pack** | 5 | 5 | 6 | 8 | 7 | 8 | 9 | 9 | 9 | 8 |
| **Taio** | **Dog** | **Pack** | 1 | 0 | 0 | 0 | 1 | 2 | 2 | 2 | 4 | 5 |
| **Sarabi** | **Dog** | **Pack** | 2 | 3 | 1 | 4 | 4 | 5 | 4 | 4 | 4 | 5 |
| **Kiano** | **Dog** | **Pack** | 0 | 0 | 0 | 1 | 1 | 0 | 1 | 1 | 0 | 1 |
| **Inaya** | **Dog** | **Pack** | 2 | 6 | 5 | 6 | 2 | 8 | 8 | 9 | 8 | 9 |
| **Kovu** | **Dog** | **Pack** | 0 | 0 | 1 | 0 | 0 | 0 | 0 | 1 | 1 | 2 |
| **Youma** | **Dog** | **Pack** | 0 | 0 | 0 | 0 | 0 | 0 | 1 | 1 | 1 | 2 |
| **Asali** | **Dog** | **Pet** | 3 | 2 | 3 | 6 | 3 | 6 | 7 | 6 | 8 | 8 |
| **Bora** | **Dog** | **Pet** | 2 | 6 | 5 | 9 | 7 | 8 | 8 | 8 | 5 | 7 |
| **Hakima** | **Dog** | **Pet** | 2 | 2 | 2 | 5 | 5 | 5 | 9 | 7 | 8 | 9 |
| **Kilio** | **Dog** | **Pet** | 4 | 7 | 3 | 4 | 4 | 2 | 2 | 5 | 6 | 7 |
| **Pepeo** | **Dog** | **Pet** | 0 | 0 | 1 | 0 | 0 | 0 | 0 | 0 | 0 | 0 |
| **Zazu** | **Dog** | **Pet** | 3 | 8 | 8 | 8 | 9 | 8 | 9 | 9 | 9 | 8 |
| **Zhenga** | **Dog** | **Pet** | 1 | 3 | 7 | 8 | 9 | 9 | 9 | 8 | 9 | 9 |
| **Koda** | **Dog** | **Pet** | 6 | 6 | 8 | 6 |  |  |  |  |  |  |
| **Freya** | **Dog** | **Pet** | 0 | 0 | 8 | 8 | 8 | 8 | 9 | 9 | 8 | 9 |

**Supplementary Table S4:** Model estimates for the relative probability to solve each apparatus by population. The rank organizes the apparatuses from **least** likely to solve (higher number) to most likely to solve (lower number). Ranks in boldface represent the instances with notorious variation between the three populations (more than two ranks).

|  | Wolf | | Pack dog | | Pet dog | |
| --- | --- | --- | --- | --- | --- | --- |
|  | **Estimate** | **Rank** | **Estimate** | **Rank** | **Estimate** | **Rank** |
| Bowl | -3,3050755 | 4 | -2,8373758 | 4 | -2,78645721 | 5 |
| Breakable | 0,5085263 | 1 | -1,3813002 | 2 | -1,68119549 | 1 |
| Door handle | -3,957924 | **6** | -4,8937384 | **9** | -4,17637603 | **9** |
| Double tray | -4,4776021 | 7 | -4,3669225 | 8 | -3,8542599 | 8 |
| Flip box | -5,0341183 | **9** | -3,3070074 | **6** | -2,04728752 | **2** |
| Push | -3,5739944 | 5 | -3,186241 | 5 | -2,99851085 | 7 |
| Rope | -2,0405346 | **3** | -3,5596465 | **7** | -2,73109211 | **4** |
| Slide door | -4,8815923 | **8** | -2,5099521 | **3** | -2,98369694 | **6** |
| Slit | -0,8491725 | 2 | -0,1076392 | 1 | -2,41629084 | 3 |

**Supplementary Table S5:** Number of apparatuses solved per animal and session. Greener colors represent a higher number, while redder ones represent a lower number.

| **Group** | **Proportion of time interacting (logit)** | **Subject** | **Proportion of time interacting (logit)** |
| --- | --- | --- | --- |
| Pack dogs | -1.751 ± 0.491 | Enzi | -4.834 ± 0.595 |
|  |  | Hiari | 0.901 ± 0.235 |
|  |  | Imara | 0.300 ± 0.186 |
|  |  | Inaya | 0.017 ± 0.294 |
|  |  | Kiano | -2.162 ± 0.219 |
|  |  | Kovu | -2.524 ± 0.304 |
|  |  | Layla | -3.815 ± 0.541 |
|  |  | Moya | 0.252 ± 0.142 |
|  |  | Panya | -3.065 ± 0.195 |
|  |  | Sarabi | -0.704 ± 0.348 |
|  |  | Taio | -1.718 ± 0.219 |
|  |  | Youma | -2.595 ± 0.484 |
|  |  | Zuri | -2.819 ± 0.268 |
| Wolves | -2.029 ± 0.652 | Amarok | 0.075 ± 0,224 |
|  |  | Chitto | -0.590 ± 0,239 |
|  |  | Etu | 0.683 ± 0,081 |
|  |  | Geronimo | -1.136 ± 0,124 |
|  |  | Kenai | -1.948 ± 0,279 |
|  |  | Nanuk | -3.690 ± 0,515 |
|  |  | Taima | -6.395 ± 0* |
|  |  | Tala | -0.028 ± 0,248 |
|  |  | Tekoa | -3.839 ± 0,704 |
|  |  | Una | -3.749 ± 0,285 |
|  |  | Yukon | -1.704 ± 0,385 |
| Pet dogs | -0.411 ± 0.515 | Asali | 0.699 ± 0.216 |
|  |  | Bora | -0.178 ± 0.183 |
|  |  | Freya | 0.106 ± 0.419 |
|  |  | Hakima | -0.358 ± 0.307 |
|  |  | Kilio | -0.769 ± 0.242 |
|  |  | Koda | 0.221 ± 0.168 |
|  |  | Pepeo | -4.338 ± 0.563 |
|  |  | Zazu | 0.679 ± 0.298 |
|  |  | Zhenga | 0.235 ± 0.184 |

* No variation, since she never approached the apparatuses.

### Original data analysis plan and reasons behind its change

Following after Rössler & Mioduszewska et al. (2020), our original data analyses plans hinged on the use of a principal component analysis (PCA) to process the data from this study (packages “FactoMineR” and “factoextra”; Lê et al., 2008; Kassandra & Mundt, 2020). For this, we used multiple variables meant to measure apparatus-directed behaviors, most of which were previously used by Rössler, Mioduszewska and colleagues. These include the latency to approach the grid of apparatuses within one body length of the subject ("latency_grid"), the number of tasks the animals interacted with ("tasks_interacted") and the number of tasks the animals did not manage to solve after interacting with them in a given session ("tasks_interacted_unsolved").

The remaining three variables were also roughly equivalent to those used by Rössler, Mioduszewska and colleagues, but with some adaptations. We decided to measure the amount of interaction with the apparatuses in terms of duration of interaction (in seconds) instead of a discrete number of “contact” events as it was done by Rössler & Mioduszewska et al. (2020). This was done because the resulting data would be more granular like this, and it would also bypass the issue of determining whether two adjacent contacts with the apparatus form part of the same “contact” event or belong to different ones. We measured this way the amount of time interacting with the baited apparatuses (“inter_baited”) and the amount of time interacted with the apparatuses the animals had already solved (“inter_solved”). These two variables, together with the amount of time spent within one body length of the subject around the grid of the apparatuses ("grid_time") were divided by the total duration of each session. This was done because not all of the sessions had the same duration, as a session would last for either 9 minutes or as much time as the animals needed to solve all 9 boxes, whichever came first. Therefore, providing durations in terms of absolute values would misrepresent the data, as the animals who solved all 9 boxes within a session would erroneously receive low values of interaction and grid times.

Unfortunately, however some issues with three variables (“tasks_interacted”, “inter_solved”, and “tasks_interacted_unsolved”) compelled us to change the statistical design used for the experiment. For the “tasks_interacted” variable, in roughly 60% of the sessions, the animals touched all boxes, which considerably reduced its predictive power. Further, in roughly 8% of the remaining sessions, the animals touched none of the boxes, which also decreased the variable’s predictive power, since this event also coincided with no interaction with the apparatuses (“inter_baited” and “inter_solved” variables) and no tasks touched but not solved (“tasks_interacted_unsolved”). Please see **Supplementary Figure S1** for the distribution of the “tasks interacted” data.

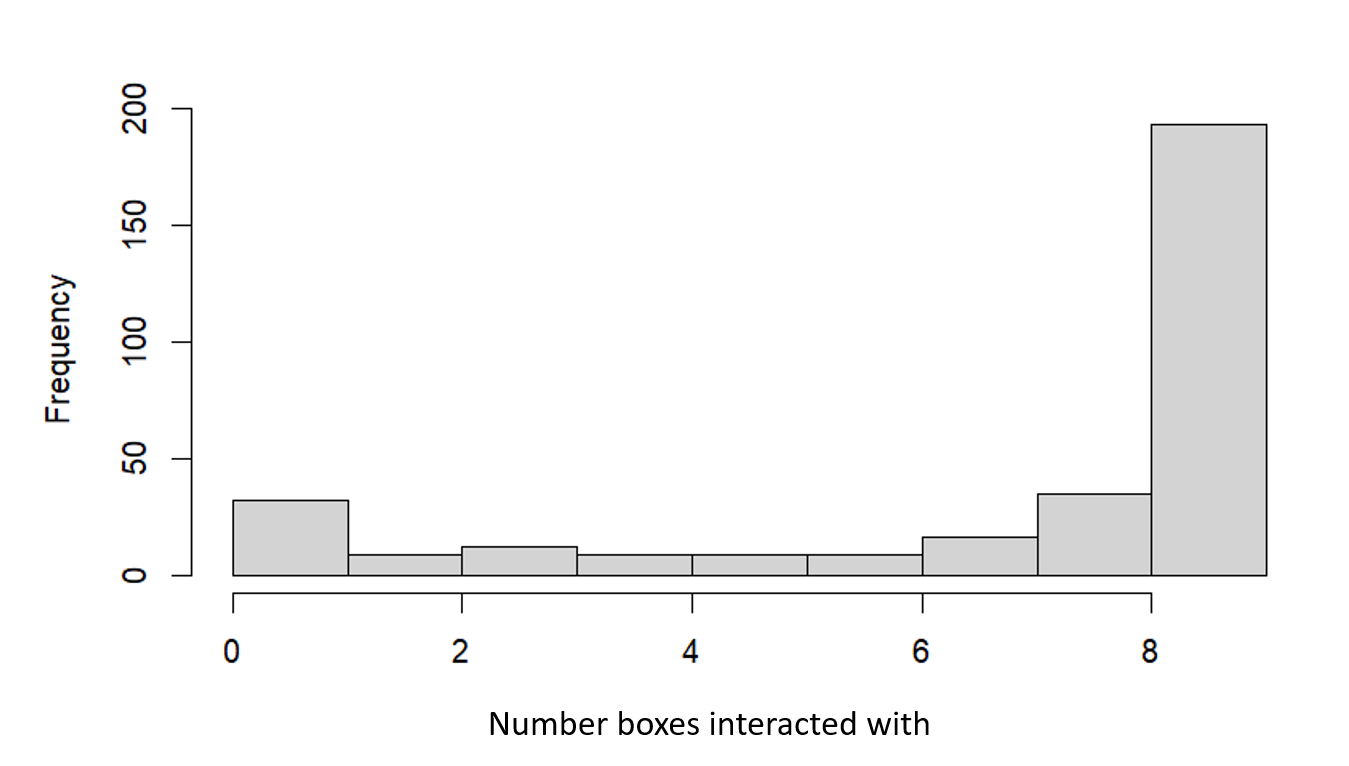

**Supplementary Figure S1:** frequency of sessions by number of boxes the animals interacted with.

The variable “inter_solved” presented another issue based on the definition of the variable and the distribution of the data. In order to interact with apparatuses that have already been solved, subjects needed to first solve at least one of the apparatuses within a given session. This greatly skewed the data, with a sizable number of sessions featuring no interaction with solved boxes (which, in roughly 95% of which, was because no apparatuses were solved), while the rest of sessions showed a normal-like distribution of the data. Please see **Supplementary Figure S2** for the distribution of the “inter_solved” data (logit-transformed).

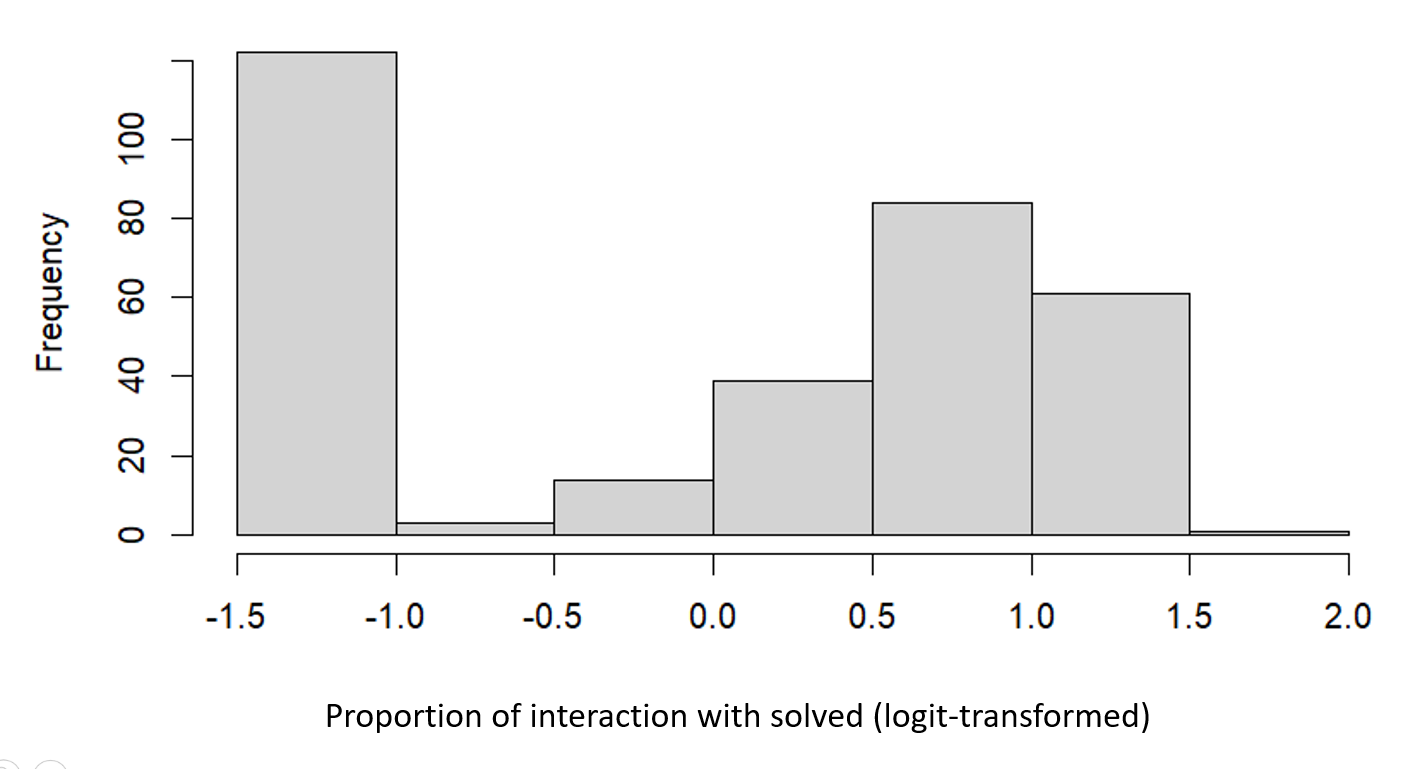
**Supplementary Figure S2:** frequency of sessions by proportion of time interacting with boxes that had already been solved (logit-transformed).

The greatest issue, however, came in the form of the “tasks_interacted_unsolved” variable. By its very nature, this variable was related to both “tasks_interacted” and the number of boxes solved, as it was, in practice, the former minus the latter. This presents several issues. The first one was that the same value of “tasks_interacted_unsolved” can be originated by several different scenarios (e.g. a value of 0 can be due to no boxes being touched, or all boxes being solved; likewise, a value of 5 could indicate 5 boxes touched but none solved, or 9 boxes touched, but 4 solved). This reduces both the predicting power of the variable and the interpretability of the results. Moreover, if animals touched all 9 boxes (as it was often the case, as stated above), this variable became, in practice, the total number of boxes minus the number of boxes solved. This made it an unsuitable variable to use in models predicting success in model solving, as it was not correlated with success in solving the apparatuses so much as an indirect measure of success in and of itself. Since this variable loaded strongly into one of the components of the PCA both in the original study and in preliminary analyses of the current one (PC2 in both), its removal had the greatest impact in the change of the data analysis design.

Due to the reasons stated above and the fact that even Rössler & Mioduszewska et al. (2020) suffered from similar problems (as corroborated by an analysis of its data, kindly provided by the authors), we decided to restructure our analyses.

Originally, we attempted to perform a PCA with the remaining variables (“latency_grid”, “grid_time”, and “inter_baited”) but, since only one of the principal components had an eigenvalue greater than 1 (our selection criterion), we decided instead to base our analyses on the most relevant variables from a biological standpoint: latency_grid (as it could signal either neophobia or neophilia) and interaction time (both with baited and solved boxes, which is a measure of persistence of exploration).
